## Supplementary material for "*CoproID* predicts the source of coprolites and paleofeces using microbiome composition and host DNA content"

#### 1 Supplementary note

##### 1.1 Preservation of coprolites

Taphonomic conditions affect the preservation potential of coprolites. These fall into two major categories: biological and sedimentological. To provide a taphonomic guide for molecular biologists working with coprolites, a reanalysis of coprolites from key sites in the desert west of North America was recently undertaken (Reinhard et al., 2019), and two common sedimentological factors affecting coprolites (even in desert environments) were identified: dry filtration of sand and precipitation of calcium-bearing moisture. Dry filtration of sand into samples can occur in horizontally deep caves, and percolation of calcium-bearing moisture is more common in open sites and at the entrances to rockshelters. Both factors can result in the complete or partial mineralization of coprolites. In dry deposits, beetles are also known to infest coprolites, and sand from the surrounding sediment matrix can become compressed into the coprolites.

Flies, beetles, mites and fungi are visible biological taphonomic agents (Reinhard et al., 2019). These decomposers are especially abundant in dense deposits of paleofeces in ancient latrines. In such latrines, the feces remain moist during the full duration of use, perhaps for decades. The communities of decomposers that establish under such conditions are strongly detrimental to the preservation of the original intestinal material. In contrast, individual fecal specimens deposited in midden (trash) contexts of short-term use often exhibit far greater preservation, especially if the fecal material is dispersed among botanical waste such as leaves, fiber masses, or cordage.

A recent review of coprolite taphonomy provides some guidelines for identifying samples with enhanced preservation potential (Reinhard et al., 2019). Flies, nematodes and especially mites are associated with poor fecal preservation at the microscopic level (Reinhard et al., 2019). These decomposers are dependent on moisture availability and microbial processes of decomposition. Visual evidence of these types of decomposers signals poor preservation potential.

Of the paleofeces analyzed in this study, only the coprolites from La Cueva de los Chiquitos Muertos, Rio Zape, Durango Mexico (Zape) have undergone a formal taphonomic assessment. From the preliminary excavation report (Brooks et al., 1962), field excavation notes were reviewed in 2019 to understand the distribution of coprolites within the cave. The Zape coprolites were dispersed in midden deposits up to 35 feet from the front of the cave. Among the midden contents, the coprolites were recovered together with the dessicated remains of maize, agave quids (expectorated agave fibers), beans, nuts, and a variety of other wild and domesticated plants (Brooks et al., 1962). The deposit covered an area 20 feet in breadth, ten feet in width and two feet in depth, and from this, it is inferred that the materials were deposited into a broad and open area within the cave. Such shallow deposits spread over a large area are known as "sheet trash." These conditions are optimal for long-term preservation, as the shallow, open-air exposure facilitates rapid drying in this arid region. The deposits, being spatially distant from the front of the cave, were protected from ambient moisture. However, they were infested with spider beetles. Such beetles prefer dry, organic substrates, and many coprolites displayed perforations left by spider beetles. It is noteworthy that these were the only decomposers identified at the site.

#### 1.2 How do we tell human from animal?

Since the 1970s, determining the biological origin of coprolites has been a focused area of research (Reinhard, 2017). Previously, only coprolites associated with skeletons or mummies could be assured a human origin, although archaeological context can also provide strong support. For example, coprolites from latrines are very likely to be human, especially when the architectural structure prohibits animal entry. However, host species determination can be less clear when coprolites are associated with trash deposits. Chame (2003) published a guide to the morphology of mammal feces and argued that scat morphology is distinct enough between species to separate human and non-human coprolites. However, dogs are clearly an exception to this system. Because of canid scatophagy, dog coprolites can closely resemble those of humans both at a macroscopic and a microscopic level.

Since the first scientific assessment of coprolites (Fry, 1977), a variety of methods have been developed to help separate human from dog samples. These methods are applied during processing and have been recently reviewed and updated (Bryant and Dean, 2006; Bryant and Reinhard, 2012; Reinhard, 2017; Reinhard and Bryant, 2008; Taylor et al., 2019). The morphology of canid coprolites is often distinct with a desiccated mucous exterior and containing evidence of hair, intact small animal bones, large splinters of long bone, cordage, cobs, pebbles, and other extraneous cave material. The rehydration stage of analysis can be insightful as researchers since Fry (1977) repeated noted that human coprolites turned the rehydration solution dark brown or opaque black, while the rehydration solution of nonhuman coprolites typically remained clear or became yellow. Most recently, Chame and her colleagues (Ferreira et al., 1989) tested feces from twenty-two South American mammalian species and found that only four species produced a dark brown or black opaque reaction upon rehydration. During the rehydration process of dog feces, the mucous also rehydrates and becomes visible on the coprolite exterior.

Reinhard and Bryant (1992) have recommended that coprolites of human origin should be based on the analysis of coprolite components. Coprolites of canine origin typically contain masses of short, nibbled dog hairs and odd inclusions, such as fragments of clothing and rope. This approach was recently augmented with the application of scanning electron microscopy (SEM; Reinhard et al. (2019)). With the aid of SEM, the details of structures such as lignin, cellulose, starch, calcium oxalate, phytoliths, and plant sporopollenin were observed, which can help identify plant remains. Finally, certain parasites are host specific and can be used to infer host species. The delicate eggs of human pinworm (Enterobius vermicularis), for example, should only be present in human coprolites, as supported by the helminth Host-Parasite Catalogue produced by the Natural History Museum, London. Since 1922, the Host Parasite Catalogue records each host-parasite record from the scientific literature, including both true infections and false infections. A review of 70,000 references (Reinhard, 2017) showed that E. vermicularis eggs had never been found in dog excreta, nor that of any non-primates. Thus, its presence in coprolite material should be considered as an additional diagnostic for human feces.

#### 2 Supplementary tables

Table S1: Host and Total read count in reference modern gut shotgun metagenomics samples

| **sample** | **host_read_count** | **all_read_count** | **percentage_host_DNA** | **organism** |
|---|---|---|---|---|
| E5 | 175600 | 11487470 | 1.53 | WHU |
| F4 | 183259 | 9696234 | 1.89 | WHU |
| E2 | 214203 | 8327324 | 2.57 | WHU |
| E1 | 227278 | 10907632 | 2.08 | WHU |
| B10 | 375059 | 11695702 | 3.21 | WHU |
| C3 | 51792 | 10324844 | 0.5 | WHU |
| B7 | 423399 | 11698532 | 3.62 | WHU |
| A1 | 153528 | 10803284 | 1.42 | WHU |
| E12 | 180817 | 12377546 | 1.46 | WHU |
| D3 | 254238 | 11408122 | 2.23 | WHU |
| C2 | 43231 | 9075014 | 0.48 | WHU |
| D7 | 225207 | 11510432 | 1.96 | WHU |
| D1 | 269617 | 12033292 | 2.24 | WHU |
| B2 | 112611 | 13075492 | 0.86 | WHU |
| A2 | 120257 | 10901438 | 1.1 | WHU |
| A6 | 160317 | 12356964 | 1.3 | WHU |
| A8 | 273006 | 10723706 | 2.55 | WHU |
| C10 | 215235 | 13017016 | 1.65 | WHU |
| C9 | 136464 | 10845174 | 1.26 | WHU |
| B8 | 340796 | 10653408 | 3.2 | WHU |
| B4 | 166426 | 10508250 | 1.58 | WHU |
| A3 | 145562 | 12421768 | 1.17 | WHU |
| D12 | 237623 | 9704844 | 2.45 | WHU |

**Table S1 continued from previous page**

| sample | host_read_count | all_read_count | percentage_host_DNA | organism |
|---|---|---|---|---|
| C11 | 284235 | 11279754 | 2.52 | WHU |
| E11 | 192428 | 11743308 | 1.64 | WHU |
| A5 | 215066 | 11882440 | 1.81 | WHU |
| B1 | 63371 | 15456544 | 0.41 | WHU |
| A11 | 199370 | 9271072 | 2.15 | WHU |
| C1 | 184712 | 11074722 | 1.67 | WHU |
| C12 | 238833 | 10917346 | 2.19 | WHU |
| D6 | 386640 | 11700012 | 3.3 | WHU |
| F1 | 262879 | 10183936 | 2.58 | WHU |
| B11 | 298888 | 10967064 | 2.73 | WHU |
| B5 | 126996 | 12667796 | 1.0 | WHU |
| D11 | 155294 | 8180156 | 1.9 | WHU |
| B3 | 133929 | 11241482 | 1.19 | WHU |
| D5 | 243972 | 11283954 | 2.16 | WHU |
| A9 | 186274 | 8874688 | 2.1 | WHU |
| C4 | 42364 | 9825904 | 0.43 | WHU |
| D2 | 177084 | 11314076 | 1.57 | WHU |
| D9 | 143234 | 10521106 | 1.36 | WHU |
| E3 | 150478 | 10696592 | 1.41 | WHU |
| E9 | 197977 | 9446418 | 2.1 | WHU |
| C7 | 180169 | 15386204 | 1.17 | WHU |
| F3 | 150425 | 11413220 | 1.32 | WHU |
| A4 | 110957 | 10818038 | 1.03 | WHU |
| E6 | 440959 | 11818510 | 3.73 | WHU |
| D8 | 196516 | 10573060 | 1.86 | WHU |
| C6 | 137728 | 14126344 | 0.97 | WHU |
| bftm2301ext33 | 55095 | 18796990 | 0.29 | NWHR |
| bftm070332 | 49029 | 26437410 | 0.19 | NWHR |
| bftm030236 | 77452 | 27571762 | 0.28 | NWHR |
| bftm2401ext31 | 7908 | 28078268 | 0.03 | NWHR |
| bftm280135 | 4046109 | 31803446 | 12.72 | NWHR |
| bftm130122 | 12870 | 41265242 | 0.03 | NWHR |
| bftm0401ecol39 | 34853 | 42149188 | 0.08 | NWHR |
| bftm230440 | 2028606 | 42879336 | 4.73 | NWHR |
| bftm1502ecol15 | 39315 | 51388140 | 0.08 | NWHR |
| bftm040212 | 11484 | 40581568 | 0.03 | NWHR |
| bftm2103ext32 | 48689 | 46270246 | 0.11 | NWHR |

**Table S1 continued from previous page**

| sample | host_read_count | all_read_count | percentage_host_DNA | organism |
|---|---|---|---|---|
| bftm140428 | 40617 | 55616436 | 0.07 | NWHR |
| bftm180134 | 21821 | 25888008 | 0.08 | NWHR |
| bftm110237 | 19889 | 43720724 | 0.05 | NWHR |
| bftm1003ext22 | 12289 | 47524028 | 0.03 | NWHR |
| bftm0803ext30 | 8469 | 18437208 | 0.05 | NWHR |
| bftm100113 | 1309446 | 100223950 | 1.31 | NWHR |
| bftm190220 | 31017 | 69358310 | 0.04 | NWHR |
| bftm270223 | 40999 | 46300184 | 0.09 | NWHR |
| bftm220210 | 26878 | 85778526 | 0.03 | NWHR |
| bftm0202ext20 | 13721 | 49753554 | 0.03 | NWHR |
| bftm0601ext18 | 7489 | 26092968 | 0.03 | NWHR |
| bftm140127 | 293440 | 104080352 | 0.28 | NWHR |
| bftm2801ext35 | 1708342 | 16746372 | 10.2 | NWHR |
| bftm1002ecol4 | 30725 | 52397704 | 0.06 | NWHR |
| bftm1303ext17 | 6928 | 28080332 | 0.02 | NWHR |
| bftm140341 | 157727 | 83257562 | 0.19 | NWHR |
| bftm2204ext27 | 367707 | 56935542 | 0.65 | NWHR |
| bftm30016 | 106606 | 74694944 | 0.14 | NWHR |
| bftm050238 | 114780 | 32861610 | 0.35 | NWHR |
| bftm1104ecol11 | 7688 | 52221098 | 0.01 | NWHR |
| bftm060125 | 20622 | 54179132 | 0.04 | NWHR |
| bftm0701ecol12 | 17077 | 39086588 | 0.04 | NWHR |
| bftm2101ext26 | 2557108 | 54682456 | 4.68 | NWHR |
| bftm2601ext24 | 143719 | 58945896 | 0.24 | NWHR |
| bftm3003ext25 | 228585 | 28016386 | 0.82 | NWHR |
| bftm09037 | 15466 | 43494168 | 0.04 | NWHR |
| bftm0604ecol14 | 26157 | 43001534 | 0.06 | NWHR |
| bftm110126 | 16884 | 37752822 | 0.04 | NWHR |
| bftm030111 | 72776 | 58457354 | 0.12 | NWHR |
| bftm290124 | 12148 | 67245490 | 0.02 | NWHR |
| bftm1704ext28 | 19128 | 48362604 | 0.04 | NWHR |
| bftm150339 | 52807 | 56123024 | 0.09 | NWHR |
| bftm2704ext23 | 194657 | 43787378 | 0.44 | NWHR |
| bftm1601ecol9 | 106670 | 73952044 | 0.14 | NWHR |
| bftm1104ext21 | 14441 | 46757422 | 0.03 | NWHR |
| bftm3004ext36 | 83706 | 30880108 | 0.27 | NWHR |
| bftm26043 | 241501 | 40648682 | 0.59 | NWHR |

**Table S1 continued from previous page**

| sample | host_read_count | all_read_count | percentage_host_DNA | organism |
|---|---|---|---|---|
| bftm2304ext34 | 409429 | 10953162 | 3.74 | NWHR |
| bftm2301ecol5 | 260661 | 74217736 | 0.35 | NWHR |
| bftm1603ecol2 | 80129 | 54867598 | 0.15 | NWHR |
| bftm120433 | 62164 | 19740812 | 0.31 | NWHR |
| bftm080321 | 19267 | 48315608 | 0.04 | NWHR |
| bftm2504ecol13 | 1344123 | 52388322 | 2.57 | NWHR |
| bftm0101ecol | 11690 | 62461836 | 0.02 | NWHR |
| bftm220116 | 160165 | 70274880 | 0.23 | NWHR |
| bftm1402ext19 | 27382 | 41748458 | 0.07 | NWHR |
| bftm17021 | 149345 | 30179114 | 0.49 | NWHR |
| bftm1701ecol6 | 11063 | 86439714 | 0.01 | NWHR |
| bftm150117 | 35070 | 77016038 | 0.05 | NWHR |
| bftm22034 | 165046 | 60784258 | 0.27 | NWHR |
| bftm0902ecol40 | 155633 | 30088172 | 0.52 | NWHR |
| bftm01028 | 10943 | 44433538 | 0.02 | NWHR |
| bftm2501ecol7 | 37260 | 53232538 | 0.07 | NWHR |
| bftm200318 | 88945 | 56072148 | 0.16 | NWHR |
| bftm080215 | 16237 | 76984636 | 0.02 | NWHR |
| bftm200129 | 212498 | 73008450 | 0.29 | NWHR |
| bftm010319 | 28985 | 63373402 | 0.05 | NWHR |
| bftm2002ext29 | 4930107 | 53649054 | 9.19 | NWHR |
| ERR1914445 | 608 | 5149226 | 0.01 | Dog |
| ERR1915141 | 199 | 3533432 | 0.01 | Dog |
| ERR1916240 | 2398 | 2959452 | 0.08 | Dog |
| ERR1915117 | 1224 | 4438944 | 0.03 | Dog |
| ERR1913401 | 2085 | 3861284 | 0.05 | Dog |
| ERR1915610 | 3612 | 3616066 | 0.1 | Dog |
| ERR1915410 | 2647 | 3802428 | 0.07 | Dog |
| ERR1916166 | 893 | 4657978 | 0.02 | Dog |
| ERR1914224 | 3162 | 7968330 | 0.04 | Dog |
| ERR1913830 | 4042 | 4956394 | 0.08 | Dog |
| ERR1915971 | 3066 | 4209920 | 0.07 | Dog |
| ERR1916218 | 865 | 4191090 | 0.02 | Dog |
| ERR1916299 | 1363 | 3491562 | 0.04 | Dog |
| ERR1915335 | 1355 | 4116570 | 0.03 | Dog |
| ERR2402762 | 18498 | 4063404 | 0.46 | Dog |
| ERR1914696 | 15239 | 4051532 | 0.38 | Dog |

Table S1 continued from previous page

| sample | host_read_count | all_read_count | percentage_host_DNA | organism |
|---|---|---|---|---|
| ERR1914207 | 7208 | 4447510 | 0.16 | Dog |
| ERR1914085 | 3503 | 4028044 | 0.09 | Dog |
| ERR1913614 | 799 | 4337814 | 0.02 | Dog |
| ERR1913472 | 37622 | 4346212 | 0.87 | Dog |
| ERR1915751 | 454 | 3857096 | 0.01 | Dog |
| ERR1916229 | 2312 | 2731206 | 0.08 | Dog |
| ERR1915582 | 253 | 4691122 | 0.01 | Dog |
| ERR1914944 | 16628 | 3581252 | 0.46 | Dog |
| ERR2402765 | 18942 | 4198878 | 0.45 | Dog |
| ERR1916302 | 1495 | 3500200 | 0.04 | Dog |
| ERR1915103 | 1230 | 4197006 | 0.03 | Dog |
| ERR1913556 | 1161 | 4701010 | 0.02 | Dog |
| ERR1914307 | 1630 | 4460470 | 0.04 | Dog |
| ERR1915539 | 3850 | 5618600 | 0.07 | Dog |
| ERR1915309 | 1978 | 4852570 | 0.04 | Dog |
| ERR1915204 | 479 | 3744596 | 0.01 | Dog |
| ERR1914185 | 6159 | 3697256 | 0.17 | Dog |
| ERR1915816 | 5834 | 3822118 | 0.15 | Dog |
| ERR1913391 | 10689 | 4830492 | 0.22 | Dog |
| ERR1914332 | 652 | 3360198 | 0.02 | Dog |
| ERR1915022 | 263 | 3385498 | 0.01 | Dog |
| ERR1914439 | 540 | 4704484 | 0.01 | Dog |
| ERR1914302 | 1499 | 3674014 | 0.04 | Dog |
| ERR1914320 | 1873 | 4628300 | 0.04 | Dog |
| ERR1913524 | 1089 | 2602736 | 0.04 | Dog |
| ERR1915317 | 1060 | 3177654 | 0.03 | Dog |
| ERR1913948 | 34470 | 5574974 | 0.62 | Dog |
| ERR1914316 | 1907 | 4707448 | 0.04 | Dog |
| ERR2402795 | 11041 | 7701058 | 0.14 | Dog |
| ERR1916189 | 403 | 4469802 | 0.01 | Dog |
| ERR1913963 | 37461 | 6034196 | 0.62 | Dog |
| ERR1915826 | 5715 | 3708276 | 0.15 | Dog |
| ERR1914111 | 2646 | 5868374 | 0.05 | Dog |
| ERR1916042 | 1166 | 4125478 | 0.03 | Dog |
| ERR1915011 | 165 | 2450928 | 0.01 | Dog |
| ERR1915380 | 1104 | 5994652 | 0.02 | Dog |
| ERR1913847 | 4226 | 5259340 | 0.08 | Dog |

**Table S1 continued from previous page**

| sample | host_read_count | all_read_count | percentage_host_DNA | organism |
|---|---|---|---|---|
| ERR1915423 | 2539 | 3681956 | 0.07 | Dog |
| ERR1913542 | 1089 | 4522812 | 0.02 | Dog |
| ERR1913392 | 2317 | 3933230 | 0.06 | Dog |
| ERR1915782 | 716 | 3017150 | 0.02 | Dog |
| ERR1915499 | 1215 | 3880756 | 0.03 | Dog |
| ERR1915611 | 3642 | 3714362 | 0.1 | Dog |
| ERR1915376 | 1025 | 5993272 | 0.02 | Dog |
| ERR1914041 | 3372 | 3524712 | 0.1 | Dog |
| ERR1914553 | 357 | 3517282 | 0.01 | Dog |
| ERR1914438 | 453 | 3627364 | 0.01 | Dog |
| ERR1916315 | 1563 | 3665318 | 0.04 | Dog |
| ERR1913675 | 1456 | 4214100 | 0.03 | Dog |
| ERR1915420 | 2338 | 3481940 | 0.07 | Dog |
| ERR1915225 | 3930 | 3020966 | 0.13 | Dog |
| ERR1914208 | 3053 | 7614094 | 0.04 | Dog |
| ERR1914674 | 1710 | 3840494 | 0.04 | Dog |
| ERR1913829 | 3673 | 4821074 | 0.08 | Dog |
| ERR1915012 | 153 | 2369958 | 0.01 | Dog |
| ERR1913953 | 35031 | 5498102 | 0.64 | Dog |
| ERR2402780 | 10542 | 7265952 | 0.15 | Dog |
| ERR1914424 | 632 | 4867382 | 0.01 | Dog |
| ERR1915578 | 259 | 4529930 | 0.01 | Dog |
| ERR1916319 | 1637 | 3652672 | 0.04 | Dog |
| ERR1915100 | 1240 | 4141642 | 0.03 | Dog |
| ERR2402772 | 18280 | 4026780 | 0.45 | Dog |
| ERR2402760 | 18150 | 4039828 | 0.45 | Dog |
| ERR1914932 | 15164 | 3287538 | 0.46 | Dog |
| ERR1913663 | 1303 | 4040726 | 0.03 | Dog |
| ERR1914441 | 668 | 4917562 | 0.01 | Dog |
| ERR1914215 | 2848 | 7596536 | 0.04 | Dog |
| ERR1915229 | 3875 | 3003368 | 0.13 | Dog |
| ERR1915197 | 445 | 3757648 | 0.01 | Dog |
| ERR1915363 | 1068 | 5700780 | 0.02 | Dog |
| ERR1916170 | 930 | 4896070 | 0.02 | Dog |
| ERR1916139 | 1827 | 4661524 | 0.04 | Dog |
| ERR1913608 | 873 | 4331384 | 0.02 | Dog |
| ERR1915231 | 4195 | 3172694 | 0.13 | Dog |

**Table S1 continued from previous page**

| sample | host_read_count | all_read_count | percentage_host_DNA | organism |
|---|---|---|---|---|
| ERR1913387 | 10563 | 4771292 | 0.22 | Dog |
| ERR1913805 | 4669 | 5005990 | 0.09 | Dog |
| ERR1916312 | 1574 | 3666764 | 0.04 | Dog |
| ERR1915253 | 888 | 2681768 | 0.03 | Dog |
| ERR1914107 | 2628 | 5828016 | 0.05 | Dog |
| ERR1916180 | 374 | 4176872 | 0.01 | Dog |
| ERR1914213 | 2976 | 7529110 | 0.04 | Dog |
| ERR1913944 | 35628 | 5713266 | 0.62 | Dog |
| ERR1914430 | 561 | 4651614 | 0.01 | Dog |
| ERR1913400 | 2197 | 3848794 | 0.06 | Dog |
| ERR1914987 | 17358 | 6702566 | 0.26 | Dog |
| ERR1914088 | 2503 | 5480602 | 0.05 | Dog |
| ERR1915765 | 577 | 3999508 | 0.01 | Dog |
| ERR1914605 | 3411 | 5270774 | 0.06 | Dog |
| ERR1913527 | 1392 | 3144084 | 0.04 | Dog |
| ERR1915320 | 1193 | 3635468 | 0.03 | Dog |
| ERR1915102 | 1360 | 4220982 | 0.03 | Dog |
| ERR1915186 | 325 | 5789872 | 0.01 | Dog |
| ERR1913947 | 34769 | 5680650 | 0.61 | Dog |
| ERR1915745 | 528 | 3839130 | 0.01 | Dog |
| ERR1915563 | 3920 | 4000478 | 0.1 | Dog |
| ERR1914040 | 3115 | 3600712 | 0.09 | Dog |
| ERR1913950 | 35171 | 5718084 | 0.62 | Dog |
| ERR1916193 | 383 | 4427438 | 0.01 | Dog |
| ERR1915453 | 342 | 3901528 | 0.01 | Dog |
| ERR1914997 | 17044 | 6507116 | 0.26 | Dog |
| ERR1914292 | 1345 | 3294266 | 0.04 | Dog |
| ERR1915122 | 216 | 3518714 | 0.01 | Dog |
| ERR1916182 | 435 | 4189138 | 0.01 | Dog |
| ERR1915558 | 3896 | 3891880 | 0.1 | Dog |
| ERR1914985 | 17576 | 6617046 | 0.27 | Dog |
| ERR1914349 | 702 | 3639940 | 0.02 | Dog |
| ERR1916187 | 471 | 4527058 | 0.01 | Dog |
| ERR1914611 | 3202 | 5193784 | 0.06 | Dog |
| ERR1915438 | 372 | 3720340 | 0.01 | Dog |
| ERR1913555 | 1133 | 4746178 | 0.02 | Dog |
| ERR1913372 | 10214 | 4489848 | 0.23 | Dog |

Table S1 continued from previous page

| sample | host_read_count | all_read_count | percentage_host_DNA | organism |
|---|---|---|---|---|
| ERR1915094 | 851 | 5057952 | 0.02 | Dog |
| ERR1915048 | 7115 | 4014060 | 0.18 | Dog |
| ERR1915455 | 402 | 3885130 | 0.01 | Dog |
| ERR1915529 | 3721 | 5474540 | 0.07 | Dog |
| ERR1913389 | 10596 | 4813572 | 0.22 | Dog |
| ERR1915571 | 4138 | 4134088 | 0.1 | Dog |
| ERR1914196 | 4277 | 2525784 | 0.17 | Dog |
| ERR1915757 | 513 | 3786956 | 0.01 | Dog |
| ERR1914667 | 1541 | 3685008 | 0.04 | Dog |
| ERR1914082 | 3314 | 3859180 | 0.09 | Dog |
| ERR1915293 | 1756 | 4254094 | 0.04 | Dog |
| ERR1915064 | 7349 | 4220568 | 0.17 | Dog |
| ERR1916078 | 1252 | 5512918 | 0.02 | Dog |
| ERR1914750 | 1693 | 4200270 | 0.04 | Dog |
| ERR1913815 | 5349 | 5771454 | 0.09 | Dog |
| ERR1913971 | 10729 | 5552042 | 0.19 | Dog |
| ERR1915838 | 5895 | 4016142 | 0.15 | Dog |
| ERR1915382 | 988 | 5989274 | 0.02 | Dog |
| ERR1913780 | 4205 | 6474574 | 0.06 | Dog |
| ERR1915590 | 261 | 4385590 | 0.01 | Dog |
| ERR1915612 | 3884 | 3611340 | 0.11 | Dog |
| ERR1915140 | 218 | 3329108 | 0.01 | Dog |
| ERR1915367 | 1013 | 5707360 | 0.02 | Dog |

Table S2: Sourcepredict source estimation of the Archaelogical samples

| | **Canis_familiaris** | **Homo_sapiens** | **Soil** | **unknown** |
|---|---|---|---|---|
| AHP003 | 0.036 | 0.017 | 0.521 | 0.426 |
| ZSM005 | 0.05 | 0.922 | 0.014 | 0.014 |
| ZSM002 | 0.05 | 0.922 | 0.014 | 0.014 |
| ZSM029 | 0.05 | 0.922 | 0.014 | 0.014 |
| BRF001 | 0.036 | 0.017 | 0.529 | 0.418 |
| ZSM025 | 0.05 | 0.922 | 0.014 | 0.014 |
| AHP002 | 0.041 | 0.019 | 0.595 | 0.345 |
| DRL001 | 0.034 | 0.016 | 0.492 | 0.458 |
| ZSM028 | 0.05 | 0.922 | 0.014 | 0.014 |
| ZSM023 | 0.05 | 0.922 | 0.014 | 0.014 |
| LEI010 | 0.036 | 0.017 | 0.524 | 0.423 |
| ECO004 | 0.034 | 0.016 | 0.491 | 0.459 |
| AHP001 | 0.032 | 0.015 | 0.469 | 0.484 |
| AHP004 | 0.527 | 0.106 | 0.009 | 0.358 |
| MLP001 | 0.031 | 0.015 | 0.455 | 0.499 |
| YRK001 | 0.949 | 0.023 | 0.014 | 0.014 |
| CMN001 | 0.033 | 0.016 | 0.486 | 0.465 |
| ZSM031 | 0.05 | 0.922 | 0.014 | 0.014 |
| ZSM027 | 0.05 | 0.922 | 0.014 | 0.014 |
| CBA001 | 0.036 | 0.017 | 0.526 | 0.421 |

Table S3: Host (nbp) and Metagenomic (Sourcepredict) compontent of the coproID prediction

| sample | nbp-human | nbp-dog | metagenomic-human | metagenomic-dog | coproID-dog | coproID_human |
|---|---|---|---|---|---|---|
| AHP001 | 0.002 | 0.998 | 0.015 | 0.032 | 0.032 | 0.0 |
| AHP002 | 0.001 | 0.999 | 0.019 | 0.041 | 0.041 | 0.0 |
| AHP003 | 0.001 | 0.999 | 0.017 | 0.036 | 0.036 | 0.0 |
| AHP004 | 0.009 | 0.991 | 0.106 | 0.527 | 0.522 | 0.001 |
| BRF001 | 0.024 | 0.976 | 0.017 | 0.036 | 0.035 | 0.0 |
| CBA001 | 0.125 | 0.875 | 0.017 | 0.036 | 0.032 | 0.002 |
| CMN001 | 0.286 | 0.714 | 0.016 | 0.033 | 0.024 | 0.004 |
| DRL001 | 0.16 | 0.84 | 0.016 | 0.034 | 0.028 | 0.003 |
| ECO004 | 0.539 | 0.461 | 0.016 | 0.034 | 0.016 | 0.008 |
| LEI010 | 0.365 | 0.635 | 0.017 | 0.036 | 0.023 | 0.006 |
| MLP001 | 0.234 | 0.766 | 0.015 | 0.031 | 0.024 | 0.003 |
| YRK001 | 0.003 | 0.997 | 0.023 | 0.949 | 0.946 | 0.0 |
| ZSM002 | 0.002 | 0.998 | 0.922 | 0.05 | 0.05 | 0.002 |
| ZSM005 | 0.87 | 0.13 | 0.922 | 0.05 | 0.007 | 0.802 |
| ZSM023 | 0.002 | 0.998 | 0.922 | 0.05 | 0.05 | 0.002 |
| ZSM025 | 0.639 | 0.361 | 0.922 | 0.05 | 0.018 | 0.589 |
| ZSM027 | 0.54 | 0.46 | 0.922 | 0.05 | 0.023 | 0.497 |
| ZSM028 | 0.544 | 0.456 | 0.922 | 0.05 | 0.023 | 0.501 |
| ZSM029 | 0.004 | 0.996 | 0.922 | 0.05 | 0.05 | 0.003 |
| ZSM031 | 0.586 | 0.414 | 0.922 | 0.05 | 0.021 | 0.54 |

Table S4: Raw sequencing read counts for Archaelogical samples

| Sample | Unique Reads | Duplicate Reads |
|---|---|---|
| ZSM002_R1 | 37111676 | 7858482 |
| ZSM002_R2 | 37248619 | 7721539 |
| ZSM005_R1 | 49004175 | 1206423 |
| ZSM005_R2 | 49069836 | 1140762 |
| ZSM023_R1 | 9823755 | 227346 |
| ZSM023_R2 | 9829938 | 221163 |
| ZSM025_R1 | 23410382 | 1481795 |
| ZSM025_R2 | 23453167 | 1439010 |
| ZSM027_R1 | 19924780 | 465551 |
| ZSM027_R2 | 19945920 | 444411 |
| ZSM028_R1 | 65372773 | 1316675 |
| ZSM028_R2 | 65465962 | 1223486 |
| ZSM029_R1 | 22163035 | 1197136 |
| ZSM029_R2 | 22183570 | 1176601 |
| ZSM031_R1 | 26261987 | 1871175 |
| ZSM031_R2 | 26300043 | 1833119 |
| AHP001_R1 | 19140976 | 365830 |
| AHP001_R2 | 19143473 | 363333 |
| AHP002_R1 | 27574180 | 569839 |
| AHP002_R2 | 27594652 | 549367 |
| AHP003_R1 | 18974980 | 360583 |
| AHP003_R2 | 18977042 | 358521 |
| AHP004_R1 | 19634617 | 361873 |
| AHP004_R2 | 19638201 | 358289 |
| BRF001S0_R1 | 9442462 | 185962 |
| BRF001S0_R2 | 9444872 | 183552 |
| CBA001S0_R1 | 8826456 | 153236 |
| CBA001S0_R2 | 8828636 | 151056 |
| CMN001_R1 | 8015084 | 127151 |
| CMN001_R2 | 8013821 | 128414 |
| DRL001S0_R1 | 7609890 | 138037 |
| DRL001S0_R2 | 7618507 | 129420 |
| ECO004_R1 | 9565407 | 158405 |
| ECO004_R2 | 9565984 | 157828 |
| LEI010S0_R1 | 821253 | 6683465 |
| LEI010S0_R2 | 1063210 | 6441508 |

**Table S4 continued from previous page**

| Sample | Unique Reads | Duplicate Reads |
|---|---|---|
| MLP001S0_R1 | 1115199 | 6208693 |
| MLP001S0_R2 | 1295090 | 6028802 |
| YRK001S0_R1 | 10830480 | 534437 |
| YRK001S0_R2 | 10845506 | 519411 |

Table S5: Modern human reference microbiome datasets for gut host DNA estimation

| Metagenome source | Food production | Other | N | Source |
|---|---|---|---|---|
| Homo sapiens - USA (Boston) | WHU | long term (>50 years) type 1 diabetes | 49 | This study |
| Homo sapiens - Burkina Faso | NWHR | - | 69 | This study |

### 3 Supplementary figures

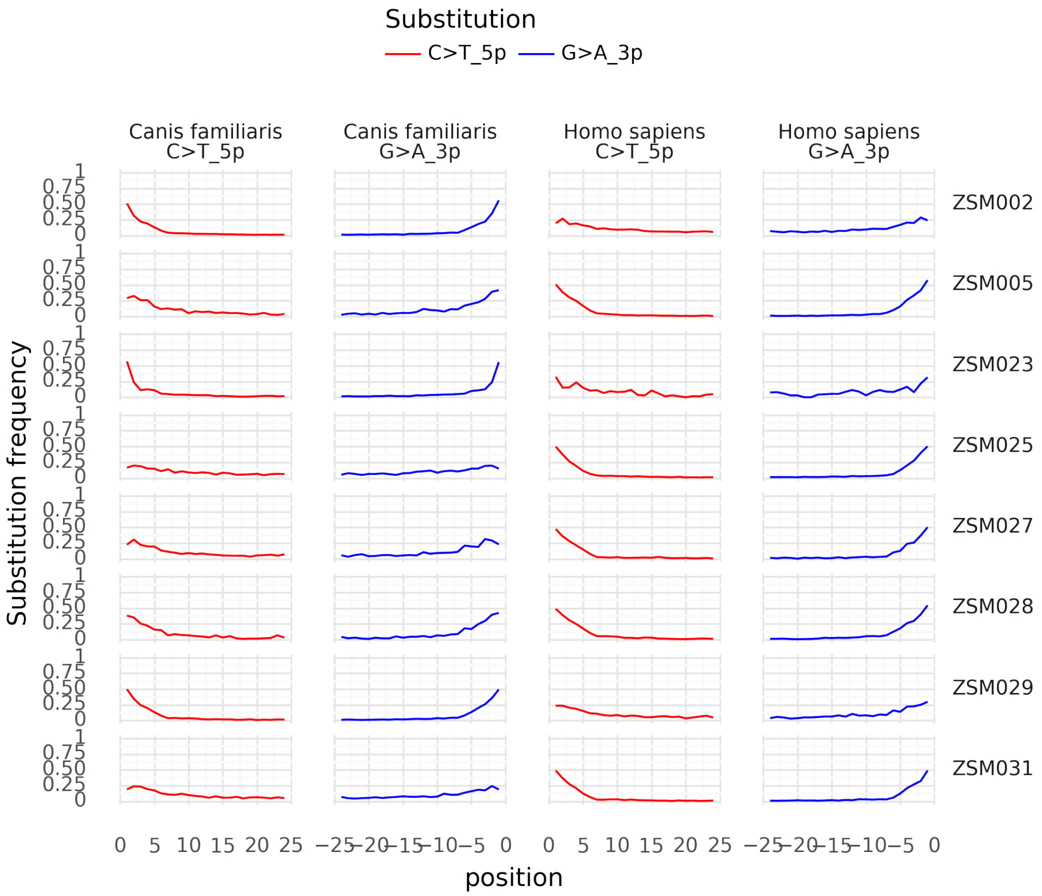


Figure S1: **Damage plot for non-UDG treated samples**

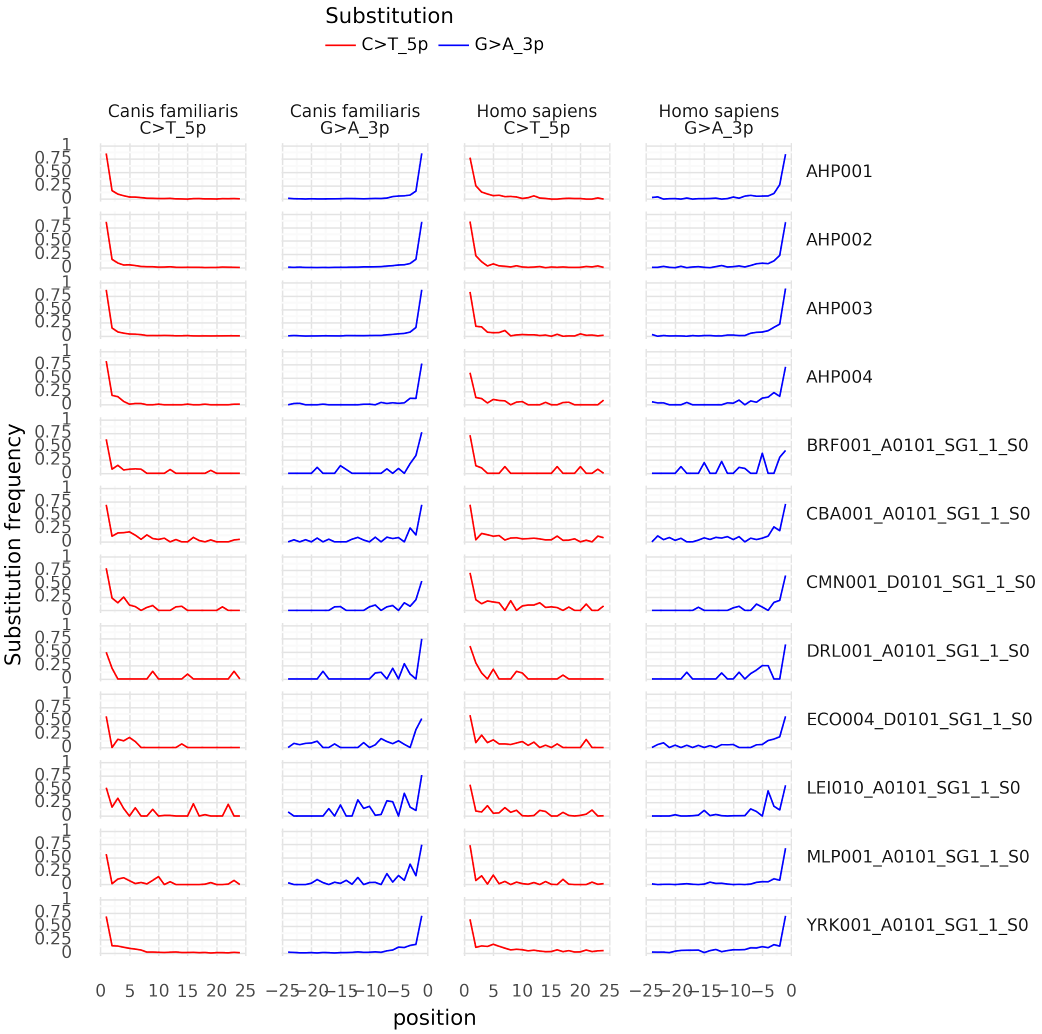


Figure S2: **Damage plot for UDG-half treated samples**
